# Cognitive Changes associated with Alzheimer’s disease in Down syndrome

**DOI:** 10.1101/263095

**Authors:** Nicholas C. Firth, Carla M. Startin, Rosalyn Hithersay, Sarah Hamburg, Peter A. Wijeratne, Kin Y. Mok, John Hardy, Daniel C. Alexander, The LonDownS Consortium, André Strydom

**Author notes:** Corresponding author details: Dr Nicholas Firth, Centre for Medical Image Computing, Department of Computer Science, University College London, London, WC1E 6BT, UK.

## Abstract

**Objective:** Individuals with Down syndrome (DS) have an extremely high genetic risk for Alzheimer’s disease (AD) however the course of cognitive decline associated with progression to dementia is ill-defined. Data-driven methods can estimate long-term trends from cross-sectional data while adjusting for variability in baseline ability, which complicates dementia assessment in those with DS.

**Methods:** We applied an event-based model to cognitive test data and informant-rated questionnaire data from 283 adults with DS (the largest study of cognitive functioning in DS to date) to estimate the sequence of cognitive decline and individuals’ disease stage.

**Results:** Decline in tests of memory, sustained attention / motor coordination, and verbal fluency occurred early, demonstrating that AD in DS follows a similar pattern of change to other forms of AD. Later decline was found for informant measures. Using the resulting staging model, we showed that adults with a clinical diagnosis of dementia and those with APOE 3:4 or 4:4 genotype were significantly more likely to be staged later, suggesting the model is valid.

**Interpretation:** Our results identify tests of memory and sustained attention may be particularly useful measures to track decline in the preclinical/prodromal stages of AD in DS whereas informant-measures may be useful in later stages (i.e. during conversion to dementia, or post-diagnosis). These results have implications for the selection of outcome measures of treatment trials to delay or prevent cognitive decline due to AD in DS. As clinical diagnoses are generally made late into AD progression, early assessment is essential.

## 1. Background

Down syndrome (DS) is due to full or partial trisomy, translocation, or mosaicism of chromosome 21, and is associated with intellectual disability (ID)^1^. DS is also a genetic cause of Alzheimer’s disease (AD), largely due to triplication of the amyloid precursor protein *(APP)* gene at 21q21.3^2^. AD neuropathology is universally present in adults with DS from their fourth decade^3,4^, driving an elevated risk for dementia due to AD that is estimated to reach 80% by age 65^5^, though the age of clinical dementia onset shows large variability^6^. The age of onset for dementia in DS is similar to that in familial AD due to mutations in the known AD causing genes - *APP,* presenilin 1 *(PSEN1)* and presenilin 2 *(PSEN2)^7^,* with mean age of diagnosis around 55, and an interquartile range of approximately 50 – 59 years of age^6^. However, unlike familial AD, the sequence and course of dementia in DS is less well described, despite this population currently accounting for the majority of genetic AD cases.

In individuals with DS, the development of dementia needs to be understood in the context of a complex cognitive phenotype that not only includes general ID, but also specific impairments in executive function, memory, language, and motor domains^8,9^. Such pre-existing impairments in those with DS need to be distinguished from subsequent decline, and in combination with varying baseline abilities and limitations in speech abilities can make the interpretation of cognitive test data, and thus clinical diagnosis, difficult^10,11^.

The evaluation of longitudinal change suggestive of AD in DS, within and across different cognitive domains, poses significant challenges. In addition to the aforementioned difficulties of assessing decline in the presence of varying degrees of premorbid ID, it is not trivial to understand the long-term longitudinal progression of a disease when the majority of studies sample populations at different stages of disease progression, by taking cross-sectional or short-term longitudinal measurements^12^. Data-driven methods have become a valuable tool for studying long-term disease progression due to their ability to estimate long-term trends from cross-sectional and short-term longitudinal snapshots of cohorts, and can be adjusted for variability in baseline ability. The event-based model (EBM) is one such method capable of estimating orderings of multimodal measurements and staging participants^13^. The EBM has been applied previously to neuroimaging, cerebrospinal fluid (CSF), and cognitive markers in sporadic AD^14^, and more recently was reformulated to model more complex cognitive datasets in young onset AD and posterior cortical atrophy^15^.

The aim of this work was to characterise the cognitive deterioration caused by AD in DS. We applied the data-driven EBM to markers of cognitive and informant-rated ability of individuals with DS to estimate the order of cognitive decline and assign participants to a disease stage. We further aimed to determine the effect of a clinical diagnosis of dementia and *APOE* genotype on stage, given that *APOE* genotype is strongly associated with age of onset of dementia due to AD, with the *e4* allele driving earlier onset and increased risk and the *e2* allele reducing risk^16^.

## 2. Methods

### 2.1. Ethics and consent

We obtained ethical approval from the National Health Service Research Ethics Committee for the LonDownS consortium’s longitudinal study of cognitive ability in DS, including approval for collection of DNA samples (13/WA/0194). Individuals with capacity to consent for themselves provided written informed consent, and for those who did not have decision-making capacity, a consultee was approached to indicate their agreement to the individual’s participation, in accordance with the UK Mental Capacity Act 2005.

### 2.2. Participants

We recruited individuals with a clinical diagnosis of DS aged 16 years and older living in family homes and residential facilities across England and Wales via a volunteer database, support groups, and local National Health Service (NHS) Trust sites. Participants with any acute physical or mental health condition were excluded from participation until they had recovered.

Due to the increased risk of people with DS developing AD neuropathology characterised by amyloid deposition beyond age 35 as demonstrated by neuropathological and amyloid positron emission tomography studies^2,17,18^, we used age 35 to split participants into two age groups. The young adult (YA) group, aged 16 – 35, were likely performing at or near to their cognitive peak, while the older adult (OA) group, aged 36 and older, were expected to have AD neuropathology with individuals presenting with varying degree of cognitive decline. We therefore defined the YA group as a pre-decline ″control″ group, while the OA group was used as a ″pathological″ group in the model.

### 2.3. Cognitive test battery

We selected tests and outcomes from the LonDownS cognitive battery that showed good psychometric properties across the age groups:

1. General cognitive abilities were assessed using raw scores from the verbal and non-verbal subscales of the Kaufman Brief Intelligence Test 2 (KBIT-2)^19^.
2. Visuospatial associate memory was assessed with the first trial memory score from the CANTAB paired associates learning (PAL) task^20^. This sums the number of pattern positions correctly recalled after their first presentation in all stages attempted.
3. Object memory was assessed using an adapted form of the Fuld object memory test^21^. This task provides measures of immediate and 5-minute delayed memory recall.
4. Orientation abilities were assessed by asking participants questions about when it was (the day, month, and year), and where they were^22^.
5. The intra/extra dimensional set shift (IED) task is a measure of rule learning and set shifting from the CANTAB^20^. Here we used the total number of stages completed.
6. An adapted version of the Tower of London for individuals with an ID assessed working memory and planning^9,23^.
7. To measure semantic verbal fluency participants were asked to name as many animals as possible in 1 minute.
8. The mean latency of responses in the simple reaction time (SRT) task from the CANTAB was used as a measure of attention and motor abilities^20,24^.
9. The finger-nose pointing test is a clinical measure of motor coordination^25^.
10. The car and motorbike score from the NEPSY-II – visuomotor precision task assesses hand-eye coordination^26^.

Informants (relatives or paid carers) completed standardized questionnaires. These included:

1. The Short Adaptive Behavior Scale (short ABS)^27^, adapted from the Adaptive Behavior Scale – Residential and Community (Part I)^28^, records participants’ everyday adaptive abilities.
2. The Dementia Questionnaire for People with Learning Disabilities (DLD) measures behaviours associated with cognitive decline in people with ID over the last two months^29^. Cognitive and social domains scores were included.
3. The Observer Memory Questionnaire (OMQ) measures individuals’ memory abilities over the last two months^30^. We developed a revised, shorter version, by selecting the most reliable items appropriate for use in adults with ID.

Further information about the LonDownS participants, cognitive assessments, and informant questionnaires can be found in (Startin et al. 2016), with a summary of tests and outcomes used in Table 1.

**Table 1.** Summary of assessments used.

|  | Test name | Primary abilities assessed | Description | Outcomes and score ranges |
| --- | --- | --- | --- | --- |
| Participant tests | Kaufmann Brief Intelligence Test 2 (KBIT-2) | General cognitive abilities | Subtests assess participants' verbal abilities (verbal knowledge and riddles) and non-verbal abilities (matrices). | Verbal raw score (0-108); Non-verbal raw score (0-46) |
|  | CANTAB paired associates learning (PAL) | Visuospatial associate memory | Participants were required to remember locations of an increasing number of patterns, hidden behind boxes on a computer screen. | First trial memory score (0-26) |
|  | Object memory test | Recall memory | Participants were required to name and remember a series of objects, then recall them in two immediate trials and one 5 minute delayed trial. | Immediate recall (0-14); Delayed recall (0-7) |
|  | CAMCOG orientation | Orientation | Assesses participants' knowledge of when it is and where they are. | Total score (0-12) |
|  | CANTAB intra/extra dimensional set shift (IED) | Rule learning and set shifting | Participants were required to learn rules about which was the 'correct' of two presented patterns on a computer screen, with a rule change after 6 consecutive correct trials. | Number of stages completed (0-9) |
|  | Tower of London | Working memory and planning | Participants were required to move beads on a board to match presented configurations. | Total score (0-10) |
|  | Verbal fluency | Semantic verbal fluency | Participants were asked to name as many animals as possible in one minute. | Number of unique animals (0-N/A) |
|  | CANTAB simple reaction time (SRT) | Attention / motor abilities | Participants were required to press a button as soon as a white square appeared on a computer screen. | Mean latency (N/A) |
|  | Finger-nose pointing | Motor coordination | Participants alternatively touch their nose and a red circle 45cm away for 20 seconds. | Total number of times the circle is touched (0-N/A) |
|  | NEPSY-II visuomotor precision | Hand-eye coordination | Participants were timed as they traced around train, car, and motorbike tracks, with times and number of errors for each track used to determine overall scores. | Car and motorbike score (0-52) |
| Informant ratings | Short Adaptive Behavior Scale (Short ABS) | Adaptive abilities | Informants answer questions about everyday adaptive abilities. | Total score (0-113) |
|  | Dementia Questionnaire for People with Learning Disabilities (DLD) | Memory and orientation / adaptive abilities | Informants answer questions about behaviours associated with cognitive decline over the last two months. | Cognitive abilities (0-44); Social abilities (0-60) |
|  | Revised Observer Memory Questionnaire (OMQ) | Memory | Informants answer questions about individuals' memory abilities over the last two months. | Total score (18-90) |

### 2.4. Imputation

Floor effects and difficulty to engage in cognitive tasks are a significant issue in studies of cognitive decline in individuals with DS, and excluding those who score at floor or who do not engage could significantly bias analyses. We therefore imputed scores as follows: individuals who attempted tasks but were clearly unable to understand task instructions were allocated a score of zero for outcomes aside from SRT mean latency, where the poorest score recorded was given. Missing items from the DLD and OMQ were imputed for up to 15% of items within each domain with the nearest integer to the mean value of completed scores.

### 2.5. Intellectual disability (ID) severity score

Premorbid ID level was defined according to the ICD10 diagnostic system, and classified into three levels based on caregiver’s reports of the individual’s best ever level of functioning – mild, moderate and severe ID, corresponding to the general functional abilities associated with IQ levels of 50 – 69, 35 – 49, and <35 respectively, as described elsewhere^31^.

### 2.6. Dementia diagnoses

Dementia was defined as the presence of an existing, independent clinical diagnosis from each individual’s clinician after comprehensive clinical assessment. Clinical diagnosis has previously been shown to be reliable^32^. None of the tests used in the EBM were used to inform diagnoses.

### 2.7. Genetic analysis

Participants’ DS status was confirmed genetically using saliva or blood samples where possible; following DNA extraction, genome-wide single nucleotide polymorphism (SNP) genotyping was performed using an Illumina OmniExpressExome array (San Diego, CA, USA) at UCL Genomics, then assembled and visually inspected in GenomeStudio to confirm the presence of an additional copy of chromosome 21, mosaicism, or translocation. APOE status was determined using a Thermo Fisher Scientific Taqman assay for SNPs rs7412 and rs429358 (Waltham, MA, USA).

### 2.8. Event-based models

Scores from the cognitive tests and informant questionnaires, controlled for ID level, were used as input in the EBM, with scores termed as ‘biomarkers’ for the remainder of this paper, to be consistent with previous descriptions of the model^13,14^. Unimodal, two-component, non-parametric mixture models were fit for each of the biomarkers, these models were then used to assign probabilities *P(x|E_i_)* and P(x|¬ *E_i_)* of a biomarker measurement, *x,* being abnormal (event *E_i_* has occured) or normal (*E_i_* has not occurred) i.e. the measurement indicating dementia or not, respectively. The OA and YA groups were used to define the initial components in each mixture model, by defining each group as a component and then fitting kernel density estimations to each group separately. The fitting procedure then uses these initial components to estimate two components corresponding to a dementia and non-dementia subpopulation. This is possible as mixture modelling is a semi-supervised method capable of learning underlying patterns in the data that correspond to dementia or not, despite being only provided with the YA and OA labels. The EBM was then used to estimate the maximum likelihood ordering of events. In this work the ordering of events corresponds to the order of decline on cognitive tests and informant questionnaires, which transition outside of a premorbid range due to cognitive decline associated with AD progression. The EBM was used as previously described^33^, briefly an event sequence *S*, was optimised using Markov chain Monte Carlo (MCMC) sampling to maximise the probability of the full set of data, *X* (all biomarkers from all individuals), given by

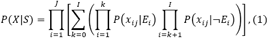

where *i∈ I* is the biomarker index and *j ∈ J* is the participant number. The fitting procedure identifies the maximum likelihood sequence *Ŝ,* from which disease stages were estimated for individuals given their test scores. Similarly to previous descriptions of the EBM (e.g.^14^), we used the stage, *k_j_*, which has the highest probability given the data and our sequence, i.e.

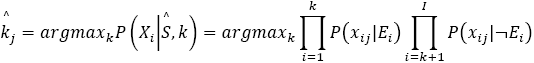

Disease stages for adults in the OA and YA groups were then compared, as were stages of adults in the OA group with and without a clinical diagnosis of dementia, and stages for individuals with different *APOE* genotypes for the YA and OA groups separately. The APOE4 group was defined as those possessing a copy of the *APOE* e4 allele *(APOE* 3:4 and 4:4), while the APOE2 group included individuals possessing a copy of the *APOE* e2 allele *(APOE* 2:2 and 2:3), and the APOE3 group consisted of those possessing two copies of the *APOE* e3 allele *(APOE* 3:3). Individuals with *APOE* 2:4 genotype were omitted from analysis. All group comparisons used Mann-Whitney U tests.

## 3. Results

This analysis included 283 participants, with details of participants included in the study summarised in Table 2.

**Table 2.**
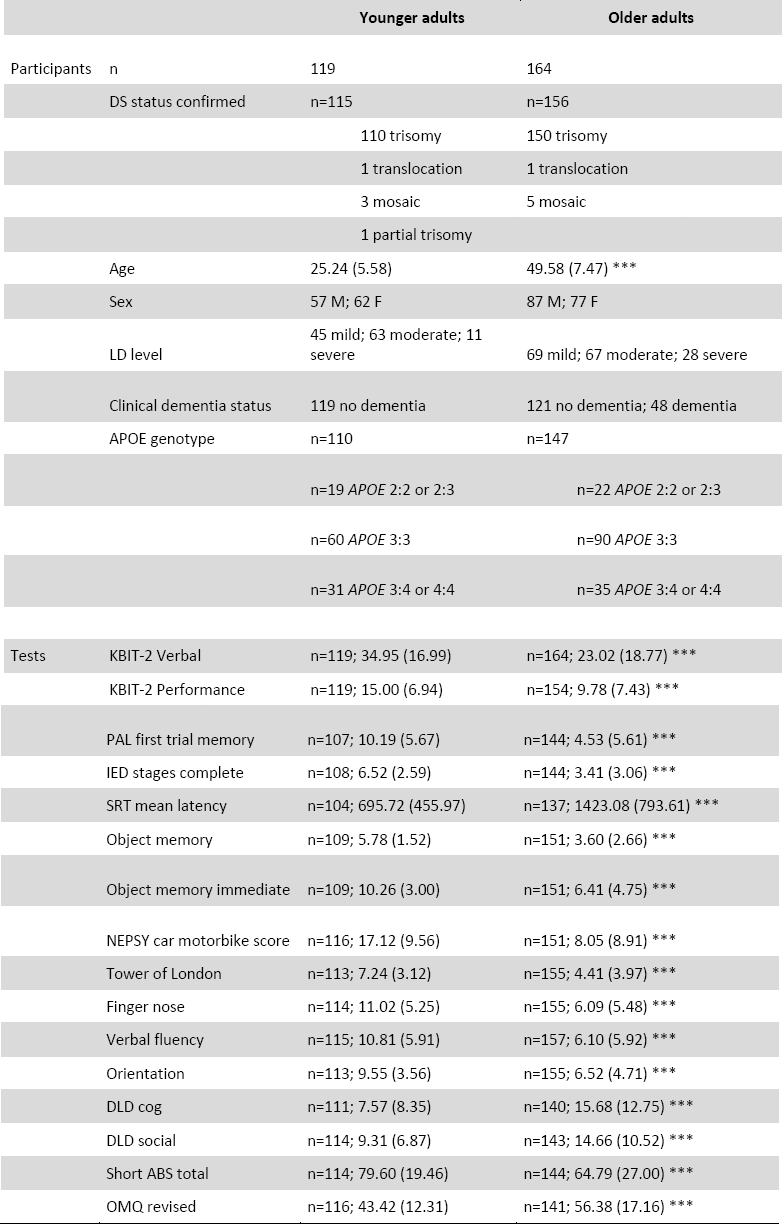
Demographic data and mean (SD) neuropsychological scores for younger adult and older adult groups. Significant differences between groups have been highlighted, p<0.05 (*), p<0.01 (**), p<0.005 (***).

|  |  | Younger adults | Older adults |
| --- | --- | --- | --- |
| Participants | n | 119 | 164 |
|  | DS status confirmed | n=115 | n=156 |
|  |  | 110 trisomy | 150 trisomy |
|  |  | 1 translocation | 1 translocation |
|  |  | 3 mosaic | 5 mosaic |
|  |  | 1 partial trisomy |  |
|  | Age | 25.24 (5.58) | 49.58 (7.47) *** |
|  | Sex | 57 M; 62 F | 87 M; 77 F |
|  | LD level | 45 mild; 63 moderate; 11 severe | 69 mild; 67 moderate; 28 severe |
|  | Clinical dementia status | 119 no dementia | 121 no dementia; 48 dementia |
|  | APOE genotype | n=110 | n=147 |
|  |  | n=19 APOE 2:2 or 2:3 | n=22 APOE 2:2 or 2:3 |
|  |  | n=60 APOE 3:3 | n=90 APOE 3:3 |
|  |  | n=31 APOE 3:4 or 4:4 | n=35 APOE 3:4 or 4:4 |
| Tests | KBIT-2 Verbal | n=119; 34.95 (16.99) | n=164; 23.02 (18.77) *** |
|  | KBIT-2 Performance | n=119; 15.00 (6.94) | n=154; 9.78 (7.43) *** |

|  |  |  |
| --- | --- | --- |
| PAL first trial memory | n=107; 10.19 (5.67) | n=144; 4.53 (5.61) *** |
| IED stages complete | n=108; 6.52 (2.59) | n=144; 3.41 (3.06) *** |
| SRT mean latency | n=104; 695.72 (455.97) | n=137; 1423.08 (793.61) *** |
| Object memory | n=109; 5.78 (1.52) | n=151; 3.60 (2.66) *** |
| Object memory immediate | n=109; 10.26 (3.00) | n=151; 6.41 (4.75) *** |
| NEPSY car motorbike score | n=116; 17.12 (9.56) | n=151; 8.05 (8.91) *** |
| Tower of London | n=113; 7.24 (3.12) | n=155; 4.41 (3.97) *** |
| Finger nose | n=114; 11.02 (5.25) | n=155; 6.09 (5.48) *** |
| Verbal fluency | n=115; 10.81 (5.91) | n=157; 6.10 (5.92) *** |
| Orientation | n=113; 9.55 (3.56) | n=155; 6.52 (4.71) *** |
| DLD cog | n=111; 7.57 (8.35) | n=140; 15.68 (12.75) *** |
| DLD social | n=114; 9.31 (6.87) | n=143; 14.66 (10.52) *** |
| Short ABS total | n=114; 79.60 (19.46) | n=144; 64.79 (27.00) *** |
| OMQ revised | n=116; 43.42 (12.31) | n=141; 56.38 (17.16) *** |

### 3.1. Event sequences

To account for the severity of cognitive deficits not caused by dementia development but instead due to the intellectual impairments associated with DS, ID level was used to estimate residuals in the YA group for each biomarker using linear regression coefficients, then these coefficients were used to calculate residuals for all individuals’ biomarker measurements. An EBM was fit using the YA and OA groups as control and disease populations respectively and a maximum likelihood event sequence was obtained together with sampling uncertainty (Figure 1a). As this method is Bayesian we did not directly estimate significance, instead a stricter estimate of uncertainty in the maximum likelihood event sequence was estimated by bootstrap resampling of the data and re-fitting the model 100 times (Figure 1b).

**Figure 1.**
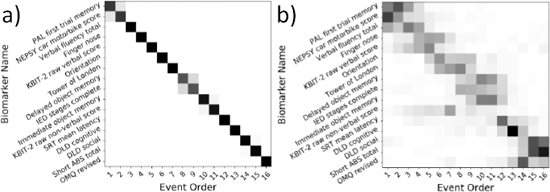
Positional variance diagrams show the maximum likelihood event sequence. Colour represents the proportion of samples with each biomarker in each position. a) Positional variance diagram of the MCMC samples generated during fitting of the EBM b) diagram of the samples generated during bootstrapping of the model.

The resulting event sequence implicates decline in visuospatial associate memory (measured using the CANTAB PAL first trial memory score), hand-eye coordination (using the NEPSY-II car/motorbike score), and semantic verbal fluency as early events associated with the likely development of significant AD neuropathology in older adults with DS. Although these tasks cover different cognitive domains, they all rely on sustained attention to perform well, indicating a common underlying ability. Tests of object memory and planning / rule learning such as the Tower of London defined mid-sequence events, while late events were defined by informant-rated scales of everyday function and cognitive ability (DLD, short ABS, and revised OMQ; Figure 1a). Bootstrapping shows a high degree of certainty in the event sequence (Figure 1b).

### 3.2. Staging

Using the maximum-likelihood event sequence a disease stage was assigned to each participant. The distribution of stages in the YA and OA groups (Figure 2) shows the YA group is significantly more likely to be at the earlier stages of the disease (U=4489.5, p<0.001). The OA group, which was assumed to show AD neuropathology, shows a spread along the event stages.

**Figure 2.**
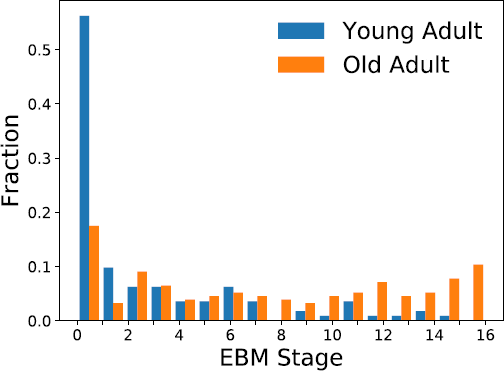
Histogram of event-based model stages for all participants, coloured by age group.

As the disease population used in this model (i.e. the OA group) consists of individuals who do not have a clinical diagnosis of dementia, we further analysed the stages of the OA group comparing those with and without clinical dementia diagnoses (Figure 3). From this we see that individuals with a clinical diagnosis of dementia are significantly more likely to be staged later according the EBM (U=743, p<0.001).

**Figure 3.**
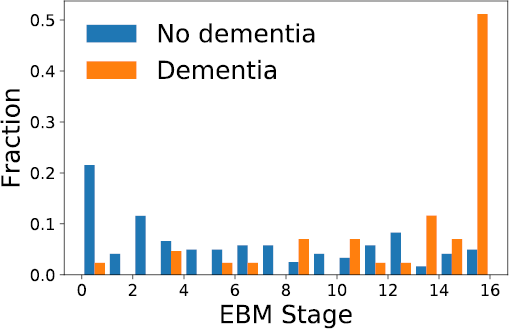
Histogram of event-based model stages for old adult participants, coloured by clinical diagnosis.

Splitting individuals based on *APOE* genotype, we saw no relationship with disease stage in the YA group (Figure 4a), however in the OA group we observed that individuals in the APOE4 group were significantly more likely to be staged later according to the EBM compared to both the APOE2 (U=205.5, p<0.002) and APOE3 (U=950, p<0.001) groups (Figure 4b).

**Figure 4.**
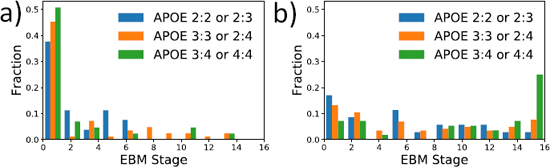
Histogram of event-based model stages coloured by APOE status. a) Young adults b) Old adults.

## 4. Discussion

Using data from nearly 300 adults with DS, we have applied an EBM to characterise the sequence of AD-related cognitive deterioration in DS. Our estimated event ordering represents the first AD progression model of this type in DS, a population at exceptionally high genetic risk of developing dementia and representing the majority of genetic AD cases. Results suggest decline in visuospatial paired associate memory, hand-eye coordination, and semantic verbal fluency may be relatively sensitive events during the prodromal stage of AD in DS. Changes in planning abilities and rule learning/shifting may occur slightly later, while changes in informant-rated behaviours and abilities appeared latest in the model. These results suggest that direct cognitive tests may be more sensitive to early changes than informant-rated questionnaires. This highlights the need for baseline cognitive assessments in this population to enable early intervention, as subtle changes in cognitive tests may be seen before carers identify decline. The staging distribution of individuals with clinical dementia diagnoses provides validity to the model, and suggests that clinical diagnoses are generally made at a relatively late stage of AD progression in DS. As additional validation of the model, we showed that older individuals with *APOE* 3:4 or 4:4 genotype were more likely to be allocated to later disease stages than those not possessing an *APOE* e4 allele.

As with sporadic AD, in DS there is a time lag of up to several decades between the development of AD neuropathology and meeting the threshold for clinical dementia diagnosis^2,34,35^. Memory decline, particularly for episodic memory, is viewed as the classic presenting symptom of AD, which gradually progresses to involve other cognitive domains^36^. However, studies of familial AD mutation carriers have found that some individuals show decline in measures of sustained attention, executive function, language, or behaviour several years prior to dementia diagnosis^37–39^, suggesting the sequence of changes we have shown in our population with DS is comparable to that seen in other genetic AD populations.

Previous EBM models have shown that in sporadic AD, changes in cognitive abilities, including memory and attention, follow changes in CSF biomarkers, but occur before brain volumetric changes^14,40^. The cognitive tests used in these models have tended to combine several cognitive processes. Here, by including tests for more specific cognitive processes as separate biomarkers, we can look more closely at the sequence of decline in populations where obtaining CSF biomarkers, for example, may be challenging.

It has been suggested that executive function decline and behavioural and personality changes may precede memory impairment in dementia development in DS^41,42^, with some studies reporting that individuals with DS may present with a frontal-like dementia syndrome in the earliest stages^41,43^. A recent systematic review of longitudinal DS studies drew similar conclusions^44^, however large variability in the follow up period and cognitive tasks used prevented a meta-analysis, and several of the studies included found, like us, that memory and spatial orientation decline seemed to happen first^45–47^.

Test sensitivity is a key challenge when assessing baseline cognition and subsequent decline in DS, and may explain these apparently conflicting findings. Some of the studies in the previously mentioned review relied on informant report, which our model suggests may be less sensitive to early change than direct cognitive assessment. Further, Lautarescu and colleagues highlighted that those with a standardized IQ < 40 had very low scores on the memory tasks used, regardless of their dementia status. The CANTAB PAL and object memory tasks used in our battery (but none of the reviewed studies) allowed us to assess visuospatial memory and immediate and delayed recall of everyday items in the majority of our sample, with fewer than 1% of our younger adults aged 16 – 35 (including many with IQ < 40) at floor on delayed object memory trials^9^, suggesting that these specific tests are suitable for those with DS and can identify decline in this population. However, for the object memory task, 40% of our younger adults were scoring at ceiling for the delayed object memory trial, suggesting this test may be insufficiently sensitive for measuring ability changes in those with comparatively strong premorbid delayed object memory. Future studies should perhaps increase the number of objects used to improve sensitivity.

The sequence of events revealed by our model has important clinical implications, and suggests that tests of visuospatial associate memory, hand-eye coordination, and verbal fluency may be particularly useful to track early, subtle cognitive change in middle-aged individuals with DS in the preclinical and prodromal stages of AD. These tests may all have decline in sustained attention in common, indicating this may be another important aspect of cognition to track changes. The model also suggests that informant-reported measures may be more useful somewhat later in the course of progression in the lead-up to dementia diagnosis, and to monitor progression after diagnosis.

AD staging based on EBM could help clinicians track decline during the early stages of cognitive decline in DS, and enable earlier diagnosis that might be beneficial by allowing for timely care-planning and support. It could also be of use to distinguish a typical sequence of events associated with AD development from the reversible decline that might be due to other, treatable, comorbidities such as depression or hypothyroidism. Equally promising is the potential use of this model to enable AD staging in clinical trials to select appropriate participants with DS for particular trials, for example those designed to prevent or delay onset of dementia in the prodromal stages. It might also be possible to use such models in the analysis of clinical trial data, by comparing different treatments or placebo controls in terms of progression along the stages of the model. However, we acknowledge that data from further longitudinal studies may be required to refine the staging model for these applications.

This analysis is based on one of the largest and most detailed studies of cognitive decline in DS to date. Participants completed a battery of cognitive tests, which were specifically designed to cover domains commonly affected by DS, including aspects of memory, motor coordination, and executive functions. The tests were adapted to be suitable for use in individuals with DS, and validated in older adults with DS^48^ before being applied in a large sample including both younger and older adults with DS which allowed for selection of tests with acceptable floor and ceiling effects^9^. By using this innovative approach and designating older adults with DS as likely being along a decline trajectory regardless of dementia status, it allowed us to provide a more complete picture of the sequence of events associated with the progression of AD in DS than has been possible to date. Furthermore, the EBM methodology allows for inclusion of individuals who had floored on some tests, which is usually a major limitation to cognitive testing in older individuals with DS^8^.

In conclusion, we have used a data driven approach to overcome some of the common issues in analysis of cognitive data in individuals with DS, and our results reveal that the sequence of events in the progression of AD in DS is comparable to events during the development of AD in other populations, including those with autosomal dominant AD. Specifically, the event sequence suggests that early decline in memory and sustained attention is followed by decline in planning and rule learning/shifting, and occurs before behavioural symptoms as reported by informants. These results help to clarify uncertainties about the sequence of events and staging of AD in DS. Future work including longitudinal data in such models will improve our understanding of decline due to AD in DS further, and will help to improve dementia diagnosis, as well as to inform selection of cognitive outcome measures in future clinical trials to prevent or delay the development of dementia during the prodromal period.

## Author contributions

AS conceived the adult cohort study in conjunction with LonDownS principal investigators. CMS, RH, and SH contributed significantly to recruitment and data collection. KYM and JH contributed to genetic analysis and interpretation of the genetic data. NF, PW, DCA, and AS designed the data analysis. NF analysed the data. NF, CMS, RH, SH, and AS wrote the paper. All authors contributed to the reviewing and editing of the paper.

## Acknowledgements

The authors would like to acknowledge all the participants in this study for their time. We would like to thank Tamara Al-Janabi for managing the project as a whole. Additional support with data collection, entry, and checking was provided by Nidhi Aggarwal, Amy Davies, Lucy Fodor-Wynne, Bryony Lowe, and Erin Rodger. This research was supported by the National Institute for Health Research networks (mental health, dementias and neurology) and participating NHS trusts. We would also like to thank our NHS network of sites that helped to identify participants. The LonDownS Consortium principal investigators are Andre Strydom (chief investigator), Department of Forensic and Neurodevelopmental Sciences, Institute of Psychiatry, Psychology and Neuroscience, King’s College London, London, UK, and Division of Psychiatry, University College London, London, UK; Elizabeth Fisher, Department of Neurodegenerative Disease, UCL Institute of Neurology, London, UK; Dean Nizetic, Blizard Institute, Barts and the London School of Medicine, Queen Mary University of London, London, UK, and Lee Kong Chian School of Medicine, Nanyang Technological University, Singapore, Singapore; John Hardy, Reta Lila Weston Institute, Institute of Neurology, University College London, London, UK, and UK Dementia Research Institute at UCL, London, UK; Victor Tybulewicz, Francis Crick Institute, London, UK, and Department of Medicine, Imperial College London, London, UK; and Annette Karmiloff-Smith, Birkbeck University of London, London, UK (deceased).

## References

1. Bittles AH, Bower C, Hussain R, Glasson EJ. The four ages of Down syndrome. Eur. J. Public Health 2007;17(2):221–225.

2. Wiseman FK, Al-Janabi T, Hardy J, et al. A genetic cause of Alzheimer disease: mechanistic insights from Down syndrome. Nat. Rev. Neurosci. 2015;16(9):564–574.

3. Mann DMA, Esiri MM. The pattern of acquisition of plaques and tangles in the brains of patients under 50 years of age with Down’s syndrome. J. Neurol. Sci. 1989;89(2- 3):169–179.

4. Wisniewski KE, Wisniewski HM, Wen GY. Occurrence of neuropathological changes and dementia of Alzheimer’s disease in Down’s syndrome. Ann. Neurol. 1985;17(3):278–282.

5. McCarron M, McCallion P, Reilly E, et al. A prospective 20-year longitudinal follow-up of dementia in persons with Down syndrome. J. Intellect. Disabil. Res. 2017;61(9):843–852.

6. Sinai A, Mokrysz C, Bernal J, et al. Predictors of Age of Diagnosis and Survival of Alzheimer’s Disease in Down Syndrome. J. Alzheimers Dis. 2018;61(2):717–728.

7. Zis P, Strydom A. Clinical aspects and biomarkers of Alzheimer’s disease in Down syndrome [Internet]. Free Radic. Biol. Med. 2017;Available from: http://www.sciencedirect.com/science/article/pii/S0891584917307402

8. Hithersay R, Hamburg S, Knight B, Strydom A. Cognitive decline and dementia in Down syndrome. Curr. Opin. Psychiatry 2017;30(2):102–107.

9. Startin CM, Hamburg S, Hithersay R, et al. The LonDownS adult cognitive assessment to study cognitive abilities and decline in Down syndrome. Wellcome Open Res. 2016;1:11.

10. Strydom A, Al-Janabi T, Houston M, Ridley J. Best practice in caring for adults with dementia and learning disabilities. Nurs. Stand. R. Coll. Nurs. G. B. 1987 2016;31(6):42–51.

11. Strydom A, Shooshtari S, Lee L, et al. Dementia in Older Adults With Intellectual Disabilities—Epidemiology, Presentation, and Diagnosis. J. Policy Pract. Intellect. Disabil. 2010;7(2):96–110.

12. Donohue MC, Jacqmin-Gadda H, Le Goff M, et al. Estimating long-term multivariate progression from short-term data. Alzheimers Dement. J. Alzheimers Assoc. 2014;10(0):S400–S410.

13. Fonteijn HM, Modat M, Clarkson MJ, et al. An event-based model for disease progression and its application in familial Alzheimer’s disease and Huntington’s disease. NeuroImage 2012;60(3):1880–1889.

14. Young AL, Oxtoby NP, Daga P, et al. A data-driven model of biomarker changes in sporadic Alzheimer’s disease. Brain 2014;137(9):2564–2577.

15. Firth NC, Primativo S, Marinescu RV, et al. Longitudinal neuroanatomical and cognitive progression of posterior cortical atrophy. submitted;

16. Patel A, Rees SD, Kelly MA, et al. Association of variants within APOE, SORL1, RUNX1, BACE1 and ALDH18A1 with dementia in Alzheimer’s disease in subjects with Down syndrome. Neurosci. Lett. 2011;487(2):144–148.

17. Annus T, Wilson LR, Hong YT, et al. The pattern of amyloid accumulation in the brains of adults with Down syndrome [Internet]. Alzheimers Dement. 2016;[cited 2015 Dec 10] Available from: http://www.sciencedirect.com/science/article/pii/S1552526015026679

18. Jennings D, Seibyl J, Sabbagh M, et al. Age dependence of brain β-amyloid deposition in Down syndrome An [18F]florbetaben PET study. Neurology 2015;84(5):500–507.

19. Kaufman AS, Kaufman NL. Kaufman Brief Intelligence Test, Second Edition. Bloomington: Pearson Inc; 2004.

20. Cambridge Cognition Ltd. CANTAB^®^. 2016;

21. Fuld PA. Guaranteed Stimulus-Processing in the Evaluation of Memory and Learning. Cortex 1980;16(2):255–271.

22. Hon J, Huppert FA, Holland A, Watson P. Neuropsychological assessment of older adults with Down’s Syndrome: An epidemiological study using the Cambridge Cognitive Examination (CAMCOG). Br. J. Clin. Psychol. 1999;38(2):155–165.

23. Strydom A, Livingston G, King M, Hassiotis A. Prevalence of dementia in intellectual disability using different diagnostic criteria. Br. J. Psychiatry 2007;191(2):150–157.

24. Edgin JO, Mason GM, Allman MJ, et al. Development and validation of the Arizona Cognitive Test Battery for Down syndrome. J. Neurodev. Disord. 2010;2(3):149–164.

25. Desrosiers J, Hébert R, Bravo G, Dutil É. Upper-extremity Motor Co-ordination of Healthy Elderly People. Age Ageing 1995;24(2):108–112.

26. Korkman M, Kirk U, Kemp S. NEPSY-II: Clinical and interpretive manual. San Antonio TX Psychol. Corp. 2007;

27. Hatton C, Emerson E, Robertson J, et al. The adaptive behavior scale-residential and community (part I): towards the development of a short form. Res. Dev. Disabil. 2001;22(4):273–288.

28. Nihira K, Leland H, Lambert NM, et al. ABS-RC:2: AAMR Adaptive Behavior Scale: residential and community. Austin, Tex.: Pro-Ed; 1993.

29. Evenhuis HM. Further evaluation of the Dementia Questionnaire for Persons with Mental Retardation (DMR). J. Intellect. Disabil. Res. 1996;40(4):369–373.

30. O’Shea MF. The cognitive and affective correlates of the memory complaint in temporal lobe epilepsy [Internet]. 1996;[cited 2017 Oct 16] Available from: http://minerva-access.unimelb.edu.au/handle/11343/35337

31. Strydom, Hassiotis A, King M, Livingston G. The relationship of dementia prevalence in older adults with intellectual disability (ID) to age and severity of ID. Psychol. Med. 2009;39(1):13.

32. Sheehan R, Sinai A, Bass N, et al. Dementia diagnostic criteria in Down syndrome. Int. J. Geriatr. Psychiatry 2015;30(8):857–863.

33. Oxtoby NP, Garbarino S, Firth NC, et al. Data Driven Sequence of Changes to Anatomical Brain Connectivity in Sporadic Alzheimer’s Disease [Internet]. Front. Neurol. 2017;8[cited 2017 Nov 6] Available from: https://www.frontiersin.org/articles/10.3389/fneur.2017.00580/abstract

34. Villemagne VL, Burnham S, Bourgeat P, et al. Amyloid β deposition, neurodegeneration, and cognitive decline in sporadic Alzheimer’s disease: a prospective cohort study. Lancet Neurol. 2013;12(4):357–367.

35. Zigman WB, Lott IT. Alzheimers disease in down syndrome: Neurobiology and risk. Ment. Retard. Dev. Disabil. Res. Rev. 2007;13(3):237–246.

36. Amieva H, Le Goff M, Millet X, et al. Prodromal Alzheimer’s disease: Successive emergence of the clinical symptoms. Ann. Neurol. 2008;64(5):492–498.

37. Fox NC, Kennedy AM, Harvey RJ, et al. Clinicopathological features of familial Alzheimer’s disease associated with the M139V mutation in the presenilin 1 gene. Pedigree but not mutation specific age at onset provides evidence for a further genetic factor. Brain 1997;120(3):491–501.

38. Ringman JM. What the Study of Persons At Risk for Familial Alzheimer’s Disease Can Tell Us About the Earliest Stages of the Disorder: A Review. J. Geriatr. Psychiatry Neurol. 2005;18(4):228–233.

39. Ryan NS, Nicholas JM, Weston PSJ, et al. Clinical phenotype and genetic associations in autosomal dominant familial Alzheimer’s disease: a case series. Lancet Neurol. 2016;15(13):1326–1335.

40. Young AL, Oxtoby NP, Huang J, et al. Multiple Orderings of Events in Disease Progression [Internet]. In: Information Processing in Medical Imaging. Springer, Cham; 2015 p. 711–722.[cited 2017 Oct 20] Available from: https://link.springer.com/chapter/10.1007/978-3-319-19992-4_56

41. Ball SL, Holland AJ, Treppner P, et al. Executive dysfunction and its association with personality and behaviour changes in the development of Alzheimer’s disease in adults with Down syndrome and mild to moderate learning disabilities. Br. J. Clin. Psychol. 2008;47(1):1–29.

42. Dekker AD, Strydom A, Coppus AMW, et al. Behavioural and psychological symptoms of dementia in Down syndrome: Early indicators of clinical Alzheimer’s disease? Cortex 2015;73(Supplement C):36–61.

43. Ball SL, Holland AJ, Hon J, et al. Personality and behaviour changes mark the early stages of Alzheimer’s disease in adults with Down’s syndrome: findings from a prospective population-based study. Int. J. Geriatr. Psychiatry 2006;21(7):661–673.

44. Lautarescu BA, Holland AJ, Zaman SH. The Early Presentation of Dementia in People with Down Syndrome: a Systematic Review of Longitudinal Studies. Neuropsychol. Rev. 2017;27(1):31–45.

45. Cosgrave MP, Tyrrell J, McCarron M, et al. A five year follow-up study of dementia in persons with Down’s syndrome: early symptoms and patterns of deterioration. Ir. J. Psychol. Med. 2000;17(1):5–11.

46. Devenny DA, Krinsky-McHale SJ, Sersen G, Silverman WP. Sequence of cognitive decline in dementia in adults with Down’s syndrome. J. Intellect. Disabil. Res. 2000;44(6):654–665.

47. Krinsky-McHale SJ, Devenny DA, Silverman WP. Changes in explicit memory associated with early dementia in adults with Down’s syndrome. J. Intellect. Disabil. Res. 2002;46(3):198–208.

48. Sinai A, Hassiotis A, Rantell K, Strydom A. Assessing Specific Cognitive Deficits Associated with Dementia in Older Adults with Down Syndrome: Use and Validity of the Arizona Cognitive Test Battery (ACTB). PLOS ONE 2016;11(5):e0153917.

